# Mechanisms of impaired cerebral hemodynamics and memory resilience differ by sex: A role for mineralocorticoid receptors in VCID

**DOI:** 10.64898/2026.10.31.763340

**Authors:** Gianna M. DeMarco, Gabrielle A. Bernstein, Dinlin Chen, James Bloete, Emily D. Schellinger, Maya E. Westbrook, Emily J. Baumoel, Jared Luchetta, Abbie C. Chapman

## Abstract

**INTRODUCTION:** Chronic hypertension is associated with cerebral hypoperfusion and is a primary risk factor for vascular contributions to cognitive impairment and dementia (VCID). Aldosterone, a key mediator of hypertension, causes cerebrovascular dysfunction via its mineralocorticoid receptor (MR).

**METHODS:** The effects of MR signaling during hypertension on hemodynamics in two cognition-centric brain regions were determined in both sexes using an established model of VCID (spontaneously hypertensive rat, SHR) and MR antagonist eplerenone. Long-term and spatial working memory were tested, and absolute brain blood flow and brain oxygenation were measured *in vivo* in CA1 of dorsal hippocampus and medial prefrontal cortex (mPFC). RESULTS: Hypertension differentially impaired hemodynamics in a sex- and region-dependent manner. Female SHR had more pervasive hypoperfusion than males that did not affect flow-metabolism coupling. Eplerenone treatment in females restored perfusion in mPFC and hippocampus, improved working memory, but did not protect hippocampal-dependent memory. Eplerenone had no effect in male SHR, indicating the MR-dependent mechanism of VCID is sex-specific.

**DISCUSSION:** This study advances our understanding of how chronic hypertension compromises brain blood flow, suggests that resilience of memory function differs by domain, and that efficacy of interventions depend upon sex.

## Introduction

There are currently over 60 million people living with dementia worldwide, and estimated to affect at least 1 in 10 people over the age of 65.^1, 2^ Vascular contributions to cognitive impairment and dementia (VCID) are the second leading cause of dementia after Alzheimer’s disease.^3^ VCID involves a spectrum of cognitive dysfunction due to cerebrovascular disease, with its most severe form being vascular dementia.^1, 4^ Importantly ∼80% of individuals clinically diagnosed with Alzheimer’s disease actually have dementia of mixed etiology – cerebrovascular pathology acting synergistically with classic Alzheimer’s disease neurodegeneration.^1, 5–7^ Thus, prevalence of VCID is likely substantially underestimated. There are currently no treatments for VCID, with therapeutic interventions focusing solely on preventative strategies mitigating risk factors. Chronic hypertension, persistently elevated blood pressure, is a leading cause of cerebrovascular disease and increases risk of VCID by 60%, specifically when hypertension develops in mid-adulthoodand.^8–11^ Chronic hypertension causes cerebrovascular dysfunction in brain regions critical to memory to contribute to VCID, including the hippocampus and prefrontal cortex.^12–17^

A critical aspect of preserving neurocognitive health is maintaining appropriate brain blood flow. In healthy adults aged 18-30 years old, perfusion in most gray matter brain regions is higher in females than males including in prefrontal cortex and hippocampus, as determined using arterial spin labeling MRI.^18, 19^ A chronic reduction in brain blood flow (hypoperfusion) occurs as a function of healthy aging; however, females undergo greater age-related hypoperfusion, specifically in the hippocampus compared to males.^20^ The hippocampus is a deep, bilateral brain structure central to learning and memory that is highly susceptible to hypoxic/ischemic injury. Chronic hypertension accelerates age-related cerebral hypoperfusion, resulting in persistent hypoxia and uncoupling of local blood flow and neuronal metabolism, ultimately leading to neurodegeneration, brain atrophy and VCID.^21, 22^ Despite the known vulnerability of the hippocampus to hypoxic/ischemic injury, how chronic hypertension influences perfusion within cognition-centric brain regions remains unclear. We recently reported that hypertension-induced cerebrovascular dysfunction occurred in a region- and sex-dependent manner.^23^ In male rats, the vasculature of the dorsal hippocampus was predominantly damaged whereas in females vascular dysfunction occurred in the prefrontal cortex.^23^ Whether hemodynamics are differentially disrupted by chronic hypertension in a sex- and brain-region specific manner remains unclear and could contribute to memory-domain specific dysfunction and VCID.

Aldosterone, a hormone produced in the kidney, is centrally involved in regulating blood pressure via the renin-angiotensin-aldosterone system. Primary aldosteronism, a disorder of pathologically high aldosterone production, is a substantial contributor to incidence of chronic hypertension.^24, 25^ Importantly, aldosterone is not only biologically active in the kidney but rather functions in a wide range of organs via interactions with the mineralocorticoid receptor (MR). MRs are expressed in the cerebrovasculature, and increased aldosterone signaling causes cerebral endothelial dysfunction and vascular remodeling during chronic hypertension in males.^26–28^ Sex steroids have a robust impact on aldosterone biosynthesis and MR signaling, including in vascular endothelium.^29–31^ Thus, understanding potential sex differences in the role of MRs in disruption of cerebral hemodynamics during chronic hypertension is critical in developing interventions to combat VCID.

In the current study, we investigated how chronic hypertension and sex impact memory functions and cerebral hemodynamics using a well-established rat model of VCID. We tested long-term and spatial working memory and measured absolute blood flow with simultaneous partial pressure of brain tissue oxygen (PbtO2) *in vivo* in two corresponding brain regions: the dorsal hippocampal CA1 subregion and medial prefrontal cortex (mPFC). The role of aldosterone-MR signaling was investigated as an underlying mechanism by which perfusion may be compromised during chronic hypertension via chronic treatment with eplerenone, a selective MR antagonist. We hypothesized that in the healthy state, females would have higher perfusion in both brain regions compared to males. We further hypothesized that chronic hypertension would differentially impair hemodynamics based upon sex, causing perfusion deficits in CA1 in males and mPFC in females, and that disruption of aldosterone signaling would protect the cerebrovasculature and improve perfusion and memory in both sexes.

## Methods

### Animals and treatments

Male and female Wistar rats and SHR were purchased from Charles River at 6 weeks of age (N=48). Animals were group housed with environmental enrichment (tunnel, chew block) on a 12-h light/dark cycle with ad libitum access to food and water in the Binghamton University - SUNY animal facility, an Association for Assessment and Accreditation of Laboratory Animal Care International (AAALAC) accredited facility. All procedures were approved by the Binghamton University – SUNY Institutional Animal Care and Use Committee (IACUC Protocol 956-25) and conducted in accordance with the NIH Guide for the Care and Use of Laboratory Animals. This study was conducted according to the ARRIVE 2.0 guidelines. Euthanasia was performed via exsanguination under chloral hydrate anesthesia followed by rapid decapitation.

Adult SHR are an established model of chronic hypertension often used to investigate hypertension-related cerebrovascular disease and cognitive decline, as they share similar neuropathology of VCID occurring in humans.^32^ Therefore, use of this model allows the investigation of how chronic hypertension impacts cerebral hemodynamics and memory functions. As uncontrolled hypertension in midlife has the greatest impact on VCID, we studied SHR at ages within mid-adulthood.^16^ Wistar-Kyoto (WKY) rats are the genetic control for SHR; however, are an established model of depression that could confound our investigation of brain blood flow as it relates to memory.^33–35^ Therefore, Wistar rats were used as normotensive controls.

SHR were randomly assigned to receive chronic treatment with selective MR antagonist eplerenone (100mg/kg/day orally in ∼ ¼ teaspoon of peanut butter; Chem Impex, Wood Dale, IL, USA) or vehicle for six weeks, beginning at ∼ 7 months old (n=8/group). Eplerenone is FDA approved for hypertension and heart failure and has fewer hormonal side-effects than spironolactone, another FDA approved MR antagonist. More specifically, eplerenone does not interfere with adrenal production of steroids such as corticosterone (cortisol in humans).^36^ The dose used has been shown to be protective of cerebrovascular function in SHR without affecting blood pressure or plasma potassium and sodium levels.^37–39^

### Behavioral Assessments of Memory Function

At ∼ 9 months of age, behavioral tasks were performed to test two types of memory function: a novel object recognition (NOR) task tested long-term memory largely dependent upon dorsal hippocampal function, and a continuous y maze task tested spatial working memory predominantly reliant on the mPFC.^14, 16^ NOR included four phases conducted over two days: 1. habituation – rats were placed in an open field arena for 5 minutes; 2. acquisition – rats explored two identical objects freely in the arena for 10 minutes; 3. consolidation – rats returned to their home cages in the animal facility and were left undisturbed for 24 hours; and 4. test – rats were placed back in the arena with one familiar and one novel object to freely explore for 5 minutes. Novel and identical object locations (left vs. right side of the arena) were randomized to control for handedness. The time spent exploring each object was quantified using automated tracking software AnyMaze 7.43 (Stoelting Co., Wood Dale, IL, USA). Recognition index was calculated as a measure of long-term memory function using the following formula: Time_Novel_ / Time_Total_, where Time_Novel_ is the time spent investigating the novel object, and Time_Total_ is the total time spent investigating both objects. Locomotion and exploratory behavior were assessed from the habituation phase of NOR (e.g., open field arena).

The day after NOR, rats were allowed to freely explore a y maze for 8 minutes. Arm entries were used to quantify an alternation index as a measure of spatial working memory using the formula: # of Alternations / (# of Arm Entries – 2). Rats were acclimated to the behavior room for one hour prior to any behavioral assessments. Male and female rats were run through behavioral paradigms on separate days to avoid potential interference, all behavioral materials/equipment were cleaned with 70% ethanol between animals, and investigators had dedicated lab coats for each sex that were laundered weekly. All behavioral assays were completed at the same time of day, ∼ 5 days prior to *in vivo* experiments.

### Assessments of Hemodynamics

To understand how chronic hypertension and sex effect hemodynamics in different brain regions, simultaneous absolute blood flow and PbtO_2_ were measured *in vivo* and compared between CA1 of dorsal hippocampus and mPFC using H_2_ clearance and an oxygen microsensor, respectively (Figure 1A and 1C), as we have previously done.^16, 40, 41^ After behavioral testing, rats were anesthetized with isoflurane (induction at 5%, maintenance during instrumentation at 2% in O_2_), intubated, and ventilated using a Rovent (Kent Scientific, Torrington, CT, USA) to control and maintain blood gases within physiological ranges. Body temperature was monitored using a rectal thermometer and maintained between 36.5 – 37.1°C with a heating pad. One femoral vein was cannulated for administration of the anesthetic chloral hydrate (50 mg/ml in sterile lactated Ringer’s solution). Both femoral arteries were cannulated to obtain continuous blood pressure recordings via a pressure transducer (TSD104A, MP200, AcqKnowledge Data Acquisition Software, BIOPAC Systems Inc., Goleta, CA, USA) and to obtain blood samples for blood gas measurements (i-STAT CG4+, Abbott, Princeton, NJ, USA). After instrumentation, rats were gradually transitioned from inhaled isoflurane anesthesia to intravenous anesthesia by tapering isoflurane down while continuously infusing chloral hydrate (30µL/min) over the course of 15 minutes. As anesthetics can have profound effects on cerebral hemodynamics, chloral hydrate was used because it is less vasoactive than isoflurane and has previously been used in studies measuring hippocampal blood flow via the hydrogen clearance method.^40, 42–44^

**Figure 1.**
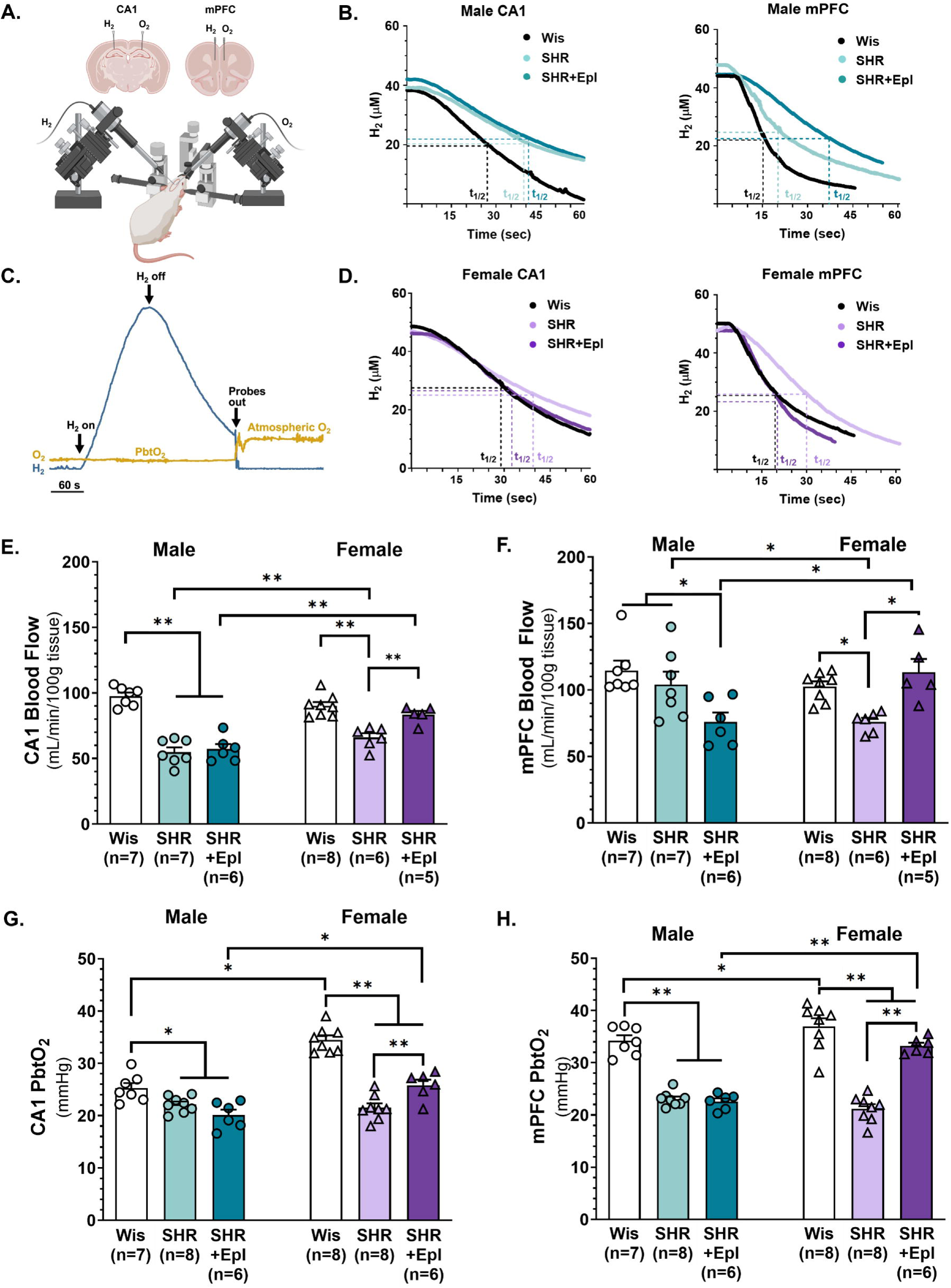
Disruption of hemodynamics in CA1 of hippocampus and medial prefrontal cortex (mPFC) during chronic hypertension. A-D. Schematic of simultaneous hydrogen (H_2_) and oxygen (O_2_) measurements in CA1 and mPFC to measure absolute blood flow and partial pressure of brain tissue O_2_ (PbtO_2_) in anesthetized rats. Representative desaturation curves from male and female normotensive Wistar rats (Wis), spontaneously hypertensive rats (SHR) and SHR chronically treated with eplerenone (SHR+Epl) demonstrate half-life of H_2_ clearance that was used to calculate rate of perfusion. **E-F.** CA1 and mPFC blood flow in males was differentially affected by chronic hypertension, showing hypoperfusion in hippocampus but not mPFC. In females, reduced perfusion occurred in both brain regions during chronic hypertension. Eplerenone treatment improved perfusion in females, however, either had no effect or decreased perfusion in males. **G-H.** PbtO_2_ was lower in CA1 and mPFC during chronic hypertension in both sexes, and was increased by eplerenone treatment only in the female sex. *p<0.05, **p<0.01; data are mean ± SEM and analyzed by two-way ANOVA with a post-hoc Tukey test.

Once under chloral hydrate anesthesia, rats were placed in prone position, secured in a dual-arm digital stereotaxic apparatus (Stoelting Co., Wood Dale, IL, USA) and the scalp retracted. Glass H_2_ and O_2_ microsensors with 50µm tips were inserted bilaterally through 2mm burr holes into CA1 of the dorsal hippocampus (from Bregma: -3.6mm posterior, 3.0mm lateral, and 2.4mm ventral at a 65° angle) or mPFC (from Bregma: +2.6mm anterior, 2.5mm lateral, and 4.5mm ventral at a 65° angle). Microsensors were calibrated daily, currents sampled at 5Hz, and data recorded using a UniAmp Multi Channel Multimeter System and data acquisition software, SensorTrace Logger (Unisense, Aarhus, Denmark). H_2_ microsensor tips contain both the sensing anode and the reference electrode (H2-50, Unisense, Aarhus, Denmark). The miniature Clark-type O_2_ microsensor tips contain a gold sensing cathode paired with a polarized, internal guard cathode and provides continuous PbtO_2_ measurements (O2-50, Unisense, Aarhus, Denmark). To measure brain blood flow, rats inhaled 4% H_2_ gas until the H_2_ current reached a steady state (saturation). After reaching saturation, H_2_ gas was turned off and tissue desaturation recorded (Figure 1B-D), as we have previously done.^16, 40^ Pilot studies were conducted to optimize the multi-site, simultaneous H_2_ and O_2_ recordings and confirm that loss of intracranial pressure over time did not impact our outcome measures. We determined that H_2_ clearance and PbtO_2_ were consistent regardless of which brain region was measured from first. Thus, bilateral burr holes were made and measurements obtained consistently in CA1 prior to mPFC.

Absolute blood flow was calculated from H_2_ desaturation curves using the initial slope to determine the first-order clearance rate in which flow (mL/sec) = 0.693/t_1/2_, where t_1/2_ is the time (sec) to reach half of the maximal tissue concentration of H_2_. The rate of clearance was then converted to mL/100g tissue/min using the following equation: blood flow = (0.693 * 100g *60sec)/ t_1/2_.^45^

Using the Fick Principle, we calculated cerebral metabolic rate of oxygen consumption (CMRO_2_; mL/min/100 g tissue) for each rat in each brain region using the formula: blood flow * (C_a_O_2_– C_v_O_2_). C_a_O_2_ is arterial oxygen content and was calculated using the following equation: C_a_O_2_ = (1.34 * Hb *SO_2_) + (0.003 * PO_2_), assuming Hb (hemoglobin) was a constant 14 g/dL in females and 15.4 g/dL in males, that SO_2_ (oxygen saturation) was a constant 0.97 (97%) and using PO_2_ as determined when blood gases were sampled. C_v_O_2_ is venous oxygen content, which we calculated using the following equation:

C_v_O_2_ = (1.34 * Hb *SO_2_) + (0.003 * PO_2_), assuming Hb was a constant as stated above, SO_2_ (oxygen saturation) was a constant 0.65 (65%) and using PO_2_ as determined when blood gases were sampled.

Brain oxygen extraction fraction (OEF; %) was calculated for each rat using the formula: CMRO_2_ / (CBF * C_a_O_2_), where CBF is cerebral blood flow as measured via H_2_ clearance.

### Estrous Staging

Estrous stages were determined in all female rats at the time of terminal experiments via vaginal cytology to understand how hormonal status may impact hemodynamics. Vaginal lavages were completed by flushing ∼200 μL of 0.1M PBS into the vaginal canal. Cell types were identified using an inverted microscope at 10X magnification and estrous stage assigned based upon the population of cells present.^23, 46, 47^

### Statistical Analyses

Sample sizes were determined by statistical power calculations using a two-sided 95% confidence interval for a single mean and 1 – β of 0.80 based on our published studies using similar methodology.^16, 48^ Statistical comparisons were made to determine main effects and interactions of sex and condition (hypertension/treatment) using a two-way ANOVA with a Tukey post-hoc test to correct for multiple comparisons. Normality of residuals was confirmed using QQ plots and Shapiro-Wilk tests. To determine differences in hemodynamics between brain regions within sex, a two-way mixed-effects model with a nested design and Tukey post-hoc test was used. Sphericity was assumed, tested with Mauchly’s Test, and Greenhouse-Geisser correction used if needed. Linear regression analyses were generated to understand the relationship between blood flow and CMRO_2_ and assess flow-metabolism coupling. Multi-linear regression analyses with Pearson’s correlations were completed and correlation matrices generated to better understand the relationship between memory function, blood pressure, blood flow and PbtO_2_. Results are reported as mean ± SEM, with p < 0.05 considered significant. All analyses were performed in GraphPad Prism 11.1. Terminal blood flow experiments were conducted in a randomized fashion (random.org). Behavioral testing was performed and analyzed by independent investigators that were blinded to group. Hemodynamic data were analyzed by two investigators that were blinded to group. Any data greater than two standard deviations from the mean were excluded from analyses, which was established *a priori*.

### Artificial Intelligence (AI) Statement

No AI tools, including generative AI, were used in any capacity in the writing of this manuscript. This includes for the construction, editing, or revising the written content, or in creation of data figures or tables.

## Results

### Chronic hypertension disrupts hemodynamics in both sexes

To understand how uncontrolled chronic hypertension in mid-adulthood impacts hemodynamics in different brain regions, and whether one sex is more susceptible to hypertension-induced hypoperfusion, absolute blood flow and PbtO_2_ were measured *in vivo* and compared between CA1 of hippocampus and mPFC in male and female SHR and normotensive Wistar rats (Figure 1A and 1C). The former is highly susceptible to hypoxic/ischemic injury and critically involved in long-term memory function, while the latter may be more sensitive to vascular damage during hypertension in females and is involved in spatial working memory.^20, 49, 50^ Table 1 shows physiological parameters of rats undergoing *in vivo* hemodynamic assessments. SHR of both sexes weighed less and had significantly higher blood pressure under chloral hydrate anesthesia compared to Wistar rats. Blood gases and blood pH were maintained within the physiological range during all hemodynamic measurements (Table 1).

**Table 1.**
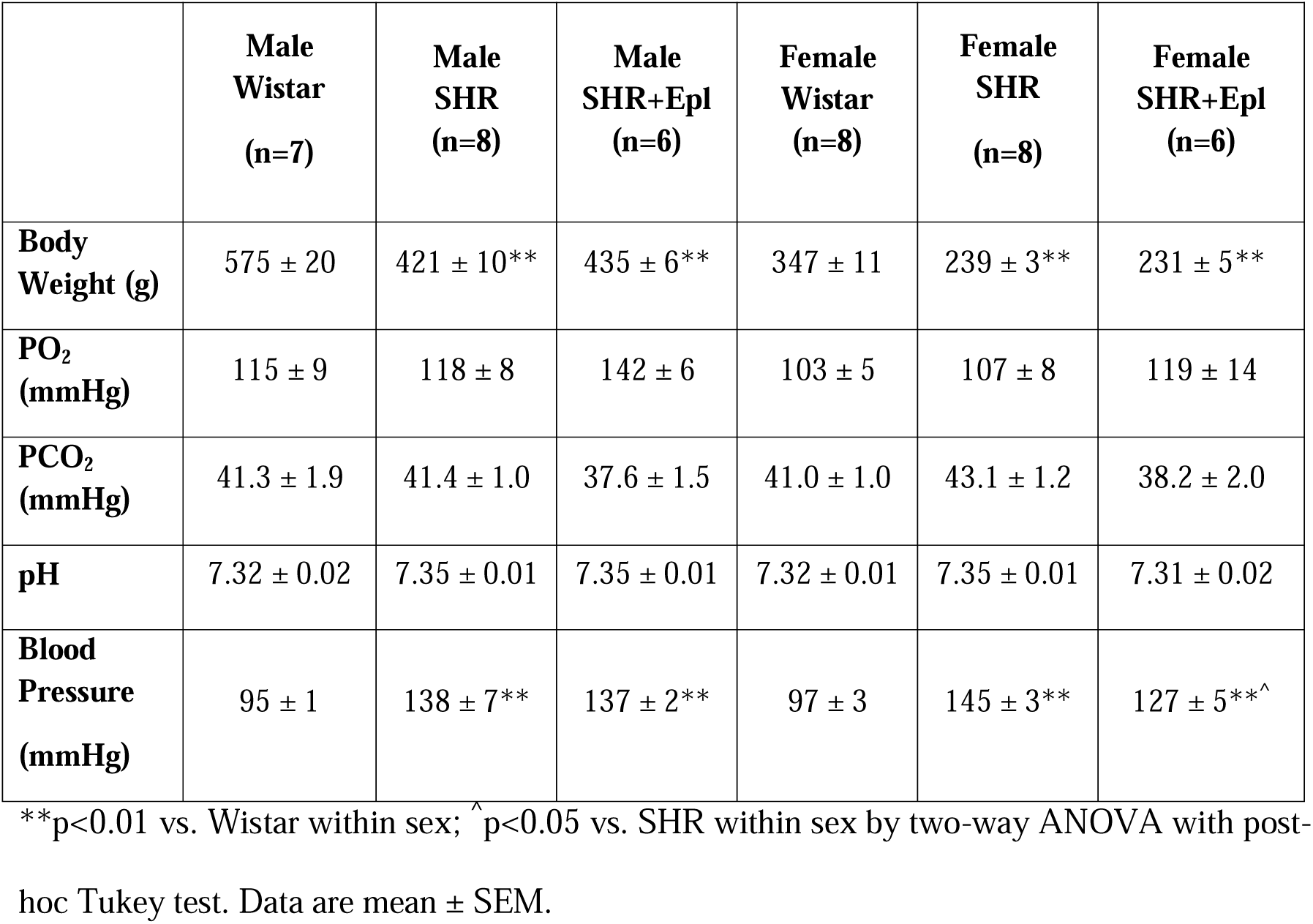
Physiological parameters under chloral hydrate anesthesia during hydrogen clearance to measure absolute blood flow and oxygen tension.

|  | **Male<br>Wistar<br>(n=7)** | **Male<br>SHR<br>(n=8)** | **Male<br>SHR+Epl<br>(n=6)** | **Female<br>Wistar<br>(n=8)** | **Female<br>SHR<br>(n=8)** | **Female<br>SHR+Epl<br>(n=6)** |
|---|---|---|---|---|---|---|
| **Body Weight (g)** | 575 ± 20 | 421 ± 10** | 435 ± 6** | 347 ± 11 | 239 ± 3** | 231 ± 5** |
| **PO<sub>2</sub> (mmHg)** | 115 ± 9 | 118 ± 8 | 142 ± 6 | 103 ± 5 | 107 ± 8 | 119 ± 14 |
| **PCO<sub>2</sub> (mmHg)** | 41.3 ± 1.9 | 41.4 ± 1.0 | 37.6 ± 1.5 | 41.0 ± 1.0 | 43.1 ± 1.2 | 38.2 ± 2.0 |
| **pH** | 7.32 ± 0.02 | 7.35 ± 0.01 | 7.35 ± 0.01 | 7.32 ± 0.01 | 7.35 ± 0.01 | 7.31 ± 0.02 |
| **Blood Pressure (mmHg)** | 95 ± 1 | 138 ± 7** | 137 ± 2** | 97 ± 3 | 145 ± 3** | 127 ± 5**<sup>^</sup> |

\*\*p<0.01 vs. Wistar within sex; <sup>^</sup>p<0.05 vs. SHR within sex by two-way ANOVA with post-hoc Tukey test. Data are mean ± SEM.

Figure 1 shows representative H_2_ desaturation curves used to calculate brain perfusion using the ½ life of clearance in each brain region in male (Figure 1B) and female rats (Figure 1D). There were main effects of sex and condition (hypertension/treatment) and a significant interaction between sex and condition on blood flow in CA1 of the hippocampus (F_1, 33_ = 13.76, p=0.0008; F_2, 33_ = 59.82, p<0.0001; F_2, 33_ = 13.29, p<0.0001 by two-way ANOVA, respectively; Figure 1E). CA1 blood flow was decreased by ∼50% in male SHR which occurred to a lesser extent in female SHR (p<0.01 by two-way ANOVA with post-hoc Tukey test) compared to normotensive controls. There were no statistical differences in CA1 blood flow between male and female normotensive rats, although females trended towards having lower CA1 perfusion levels than males (p=0.09 by two-way ANOVA with post-hoc Tukey test; Figure 1E). Oxygen tension in the hippocampus was affected by chronic hypertension similarly to CA1 blood flow. There were main effects of sex, condition, and a sex-by-condition interaction on PbtO_2_ (F_1, 37_ = 43.06, p<0.0001; F_2, 37_ = 50.56, p<0.0001; F_2, 37_ = 17.41, p<0.0001 by two-way ANOVA, respectively; Figure 1G). Under normotensive conditions, CA1 PbtO_2_ was higher in females than males and was significantly lower during chronic hypertension in both sexes (p<0.05 by two-way ANOVA with post-hoc Tukey test).

In mPFC, there was a main effect of condition, but not sex, and a significant sex-by-condition interaction on blood flow (F_2, 33_ = 4.043, p=0.0269; F_1, 33_ = 0.0247, p=0.8761; F_2, 33_ = 10.41, p=0.0003 by two-way ANOVA, respectively; Figure 1F). Blood flow in mPFC was similar between male SHR and Wistar, however, flow was significantly lower in female SHR (p<0.05 by two-way ANOVA with post-hoc Tukey test). There were no differences in mPFC blood flow between male and female normotensive rats (Figure 1F). There were main effects of sex, condition and a significant interaction between the two on mPFC PbtO_2_ (F_1, 38_ = 23.89, p<0.0001; F_2, 38_ = 104.2, p<0.0001; F_2, 38_ = 21.10, p<0.0001 by two-way ANOVA, respectively; Figure 1H). Under normotensive conditions, mPFC PbtO_2_ was higher in females than males and was significantly decreased in male and female SHR (p<0.05 by two-way ANOVA with post-hoc Tukey test). One male Wistar was excluded from blood flow and PbtO_2_ data due to excessive blood loss. Two female and one male SHR were excluded from blood flow data due to inaccurate H2 clearance measurements from improper calibrations or being a statistical outlier (>2 SD from the mean). These data suggest that chronic hypertension caused more pervasive hypoperfusion in females than males and that impact was present across both brain regions studied. Additionally, tissue oxygenation of hippocampus and mPFC was reduced during hypertension, even in the absence of perfusion deficits.

To understand potential regional differences in hemodynamics, we compared CA1 and mPFC blood flow and PbtO_2_ in male and female rats. In males, there were main effects of condition and brain region and a significant interaction between the two on blood flow (F_2, 17_ = 18.05, p<0.0001; F_1, 17_ = 34.22, p<0.0001; F_2, 17_ = 4.773, p=0.0226 by two-way mixed-effects analysis, respectively) and PbtO_2_ (F_2, 36_ = 63.71, p<0.0001; F_1, 36_ = 39.64, p<0.0001; F_2, 36_ = 15.10, p<0.0001by two-way mixed-effects analysis, respectively). Blood flow was higher in mPFC compared to CA1 in male Wistar and SHR. PbtO_2_ was higher in mPFC compared to CA1 in Wistars, but similar between regions in SHR (p<0.05 by two-way mixed-effects analysis with post-hoc Tukey test). In females, there were main effects of condition and brain region on blood flow, and a condition-by-brain region interaction that trended towards significance (F_2, 18_ = 18.44, p<0.0001; F_1, 14_ = 28.98, p<0.0001; F_2, 14_ = 3.511, p=0.0581 by two-way mixed-effects analysis, respectively). There were main effects of condition and brain region, and a significant interaction between the two on PbtO_2_ (F_2, 20_ = 76.61, p<0.0001; F_1, 19_ = 23.95, p=0.0001; F_2, 19_ = 11.40, p=0.0006 by two-way mixed-effects analysis, respectively). Similar to normotensive males, blood flow and PbtO_2_ were higher in mPFC than in CA1 in female Wistar rats, however, perfusion and oxygen levels were similar between regions in female SHR (p<0.05 by two-way mixed-effects analysis with post-hoc Tukey test). The majority of female rats were in met/diestrus (16/24, 66.7%), with fewer rats being in estrus (5/25, 20.8%) and proestrus (3/24, 12.5%). There were no detectable differences in blood flow, PbtO2 or blood pressure between estrous stages (data not shown). These findings suggest that baseline hemodynamics differ between cortex and hippocampus similarly in both sexes, and perfusion and oxygenation are compromised similarly in both regions during chronic hypertension.

### MR antagonism rescued hypertension-induced perfusion deficits only in females

Male and female SHR were chronically with the selective MR antagonist eplerenone for six weeks prior to hemodynamic assessments to understand underlying mechanisms by which perfusion may be disrupted during chronic hypertension. Eplerenone treatment did not affect body weight in either sex, and blood gases were within physiological range during cerebral hemodynamics measurements (Table 1). Blood pressure under chloral hydrate anesthesia was similar between male SHR regardless of eplerenone treatment, both of which were higher than Wistar rats. However, eplerenone treatment had a moderate anti-hypertensive effect in female SHR. Blood pressure of eplerenone-treated female SHR was ∼ 20 mmHg lower than vehicle-treated SHR, though both groups had higher blood pressure than female Wistar rats (Table 1).

MR antagonism did not affect CA1 blood flow or PbtO_2_ in male SHR. However, eplerenone treatment in female SHR increased both perfusion and PbtO_2_ in CA1 of hippocampus (p<0.05 by two-way ANOVA with post-hoc Tukey test; Figure 1E and 1G). Interestingly, mPFC blood flow was substantially decreased in male SHR treated with eplerenone compared to vehicle-treated males, but eplerenone did not further reduce PbtO_2_ (p<0.05 by two-way ANOVA with post-hoc Tukey test; Figure 1F and 1H). In females, eplerenone treatment restored both mPFC blood flow and PbtO_2_ levels to be similar to normotensive females (p<0.05 by two-way ANOVA with post-hoc Tukey test; Figure 1E and 1G). Two male and one female SHR treated with eplerenone were excluded from blood flow and PbtO_2_ data due to hypotension from excessive blood loss due to surgical complications. One female SHR treated with eplerenone was excluded from blood flow and PbtO_2_ data due to blood gases being outside the physiological ranges, and one from blood flow data for being a statistical outlier (> 2SD from the mean). These data suggest a role of MR in hypertension-induced hypoperfusion and decreased oxygenation in both hippocampus and mPFC in females, whereas MR signaling may have a protective role on perfusion during hypertension in males, specifically within the mPFC.

### Neuronal metabolism differed by sex during hypertension in a brain-region-dependent manner

Chronic hypertension is known to contribute to age-related cognitive decline, which is thought to be primarily due to reductions in basal or activity-dependent changes in blood flow.^12^ However, chronic hypertension may directly impact neuronal health in cognition-centric brain regions that could contribute to VCID. To better understand local hemodynamics, hypertension-induced changes in neuronal metabolism, and whether one sex and/or brain region was more susceptible to dysfunction that may be rescued by MR antagonism, we compared CMRO_2_ between male and female SHR and Wistar rats in CA1 of hippocampus and mPFC. There was a main effect of condition, but not sex, and a significant interaction between condition and sex on CMRO_2_ in CA1 of dorsal hippocampus (F_2,33_ = 56.52, p<0.0001; F_1, 33_ = 0.8126, p=0.3739; F_2, 33_ = 13.88, p<0.0001 by two-way ANOVA, respectively; Figure 2A). Compared to normotensive males, female Wistar rats had lower CMRO_2_ in CA1. CMRO_2_ was significantly lower in SHR compared to Wistar rats in both sexes, and eplerenone treatment restored metabolic function in female SHR to be similar to Wistar rats but had no effect in males (p<0.01 by two-way ANOVA with post-hoc Tukey test). These findings suggest CA1 neuronal dysfunction occurred during chronic hypertension in both sexes and was improved with MR blockade only in the female sex.

**Figure 2.**
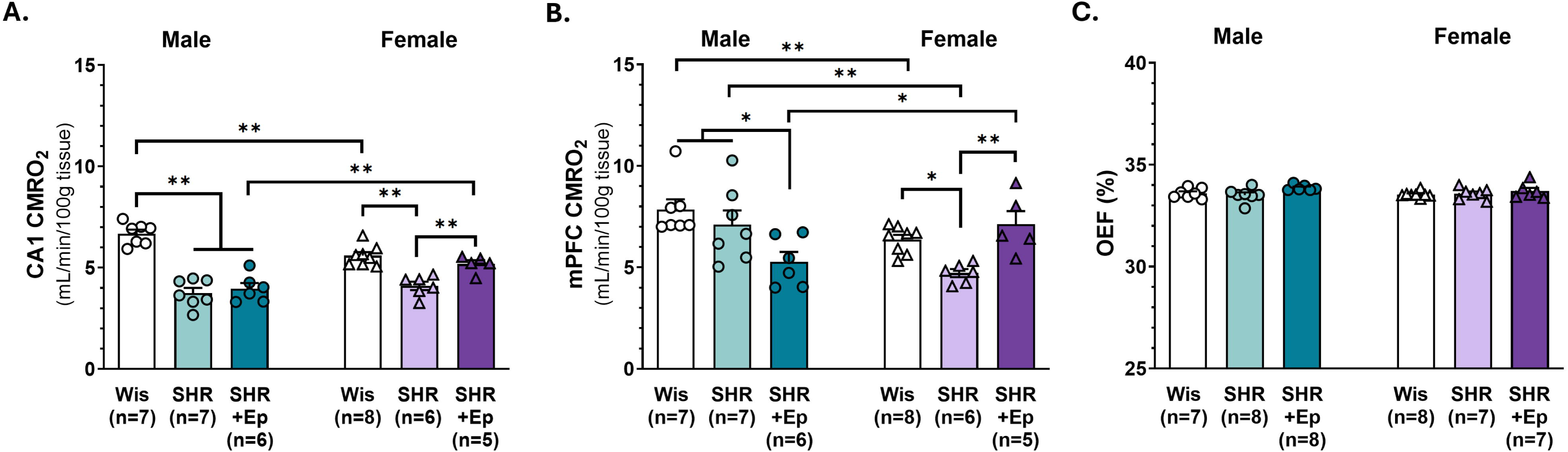
Decreased neuronal metabolism during chronic hypertension. **A.** Cerebral metabolic rate of oxygen consumption (CMRO_2_) in male and female spontaneously hypertensive rats (SHR) was lower in CA1 of the hippocampus compared to normotensive Wistar rats (Wis) that was improved in female SHR chronically treated with eplerenone (SHR+Epl) but not males. **B.** CMRO_2_ in medial prefrontal cortex (mPFC) was unaffected by chronic hypertension in males and was decreased by eplerenone treatment. In females, CMRO_2_ was lower during chronic hypertension that was restored by eplerenone treatment. **C.** Brain oxygen extraction fraction (OEF) was similar between groups. *p<0.05, **p<0.01; data are mean ± SEM and analyzed by two-way ANOVA with a post-hoc Tukey test.

In mPFC, there was a main effect of condition, but not sex, and a condition-by-sex interaction on CMRO_2_ (F_2,33_ = 3.502, p=0.0418; F_1, 33_ = 2.744, p=0.1071; F_2, 33_ = 9.303, p=0.0006 by two-way ANOVA, respectively; Figure 2B). Similar to what was found in CA1, CMRO_2_ in normotensive Wistar rats was lower in females compared to males. Chronic hypertension did not affect neuronal metabolism in male SHR, but CMRO_2_ was significantly lower in female SHR and was restored by eplerenone treatment. Interestingly, eplerenone treatment decreased CMRO_2_ in males compared to SHR and Wistar rats (p<0.01 by two-way ANOVA with post-hoc Tukey test). There were no main effects of sex or condition and no interaction between the two on brain OEF (F_1,35_ = 0.3853, p=0.5388; F_2,35_ = 3.185, p=0.0536; F_2,35_ = 0.6226, p=0.5424 by two-way ANOVA, respectively; Figure 2C). These findings suggest a sex-specific effect of chronic hypertension on neuronal dysfunction in mPFC that was due to aldosterone-MR signaling.

An essential aspect of neurovascular function critical to maintaing neuronal health is the coupling of blood flow to neuronal metabolic demand (e.g., flow-metabolism coupling). Linear regression analyses were used to determine the relationship between absolute blood flow and CMRO_2_ in CA1 and mPFC within each sex (Figure 3A and 3D) and separated by condition (hypertension/treatment; Figure 3B-C and 3E-F). There was a significant positive relationship between blood flow and CMRO_2_ that was nearly linear in male and female rats in both CA1 (Figure 3A) and mPFC (Figure 3D) that was not different between sexes. When stratified by treatment within sex, there were no differences in correlations between blood flow and metabolism in CA1 between groups of males (Figure 3B) or females (Figure 3C), nor were there differences in correlations between flow and metabolism in mPFC of males (Figure 3E) or females (Figure 3F). These data suggest that tight coupling between neuronal metabolism and perfusion was maintained during chronic hypertension in both sexes.

**Figure 3.**
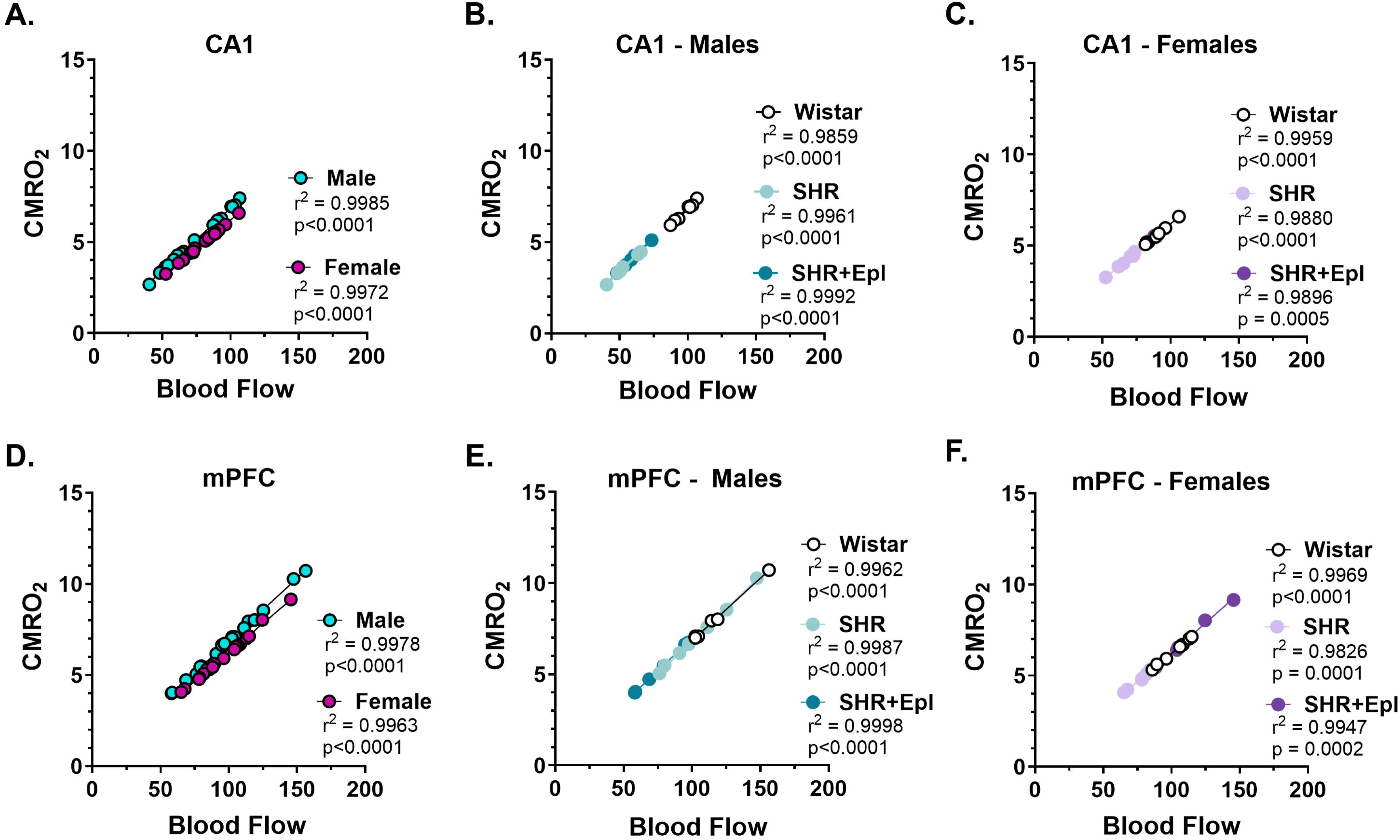
Metabolism-flow coupling was sustained in CA1 of hippocampus and medial prefrontal cortex (mPFC) during chronic hypertension in both sexes. A-C. CA1 blood flow and cerebral metabolic rate of oxygen consumption (CMRO_2_) were tightly correlated in both males and females, and unaffected in spontaneously hypertensive rats (SHR) without and with eplerenone treatment (SHR+Epl). **D-F.** In mPFC, blood flow and CMRO_2_ were correlated similarly in both sexes and were unchanged in SHR and SHR+Epl compared to normotensive Wistar rats. Data were analyzed using linear regression analyses to determine if slopes were significantly non-zero and for goodness of fit (r^2^) and are reported for each comparison.

### MR-dependent memory dysfunction and resilience during chronic hypertension is sex- and domain-dependent

To understand the potential impact of hemodynamic disruptions on memory function during chronic hypertension, and whether this differed by sex or brain region/memory domain, we used behavioral assessments of long-term memory (NOR) and spatial working memory (y maze). There were main effects of sex and condition (hypertension/treatment), but no interaction between sex and condition for locomotion, including average speed (F_1, 42_ = 88.04, p<0.01; F_2, 42_ = 7.417, p=0.0017; F_2, 42_ = 0.3659, p=0.6957 by two-way ANOVA, respectively; Figure 4A) and distance traveled (F_1, 42_ = 87.22, p<0.01; F_2, 42_ = 5.668, p=0.0066; F_2, 42_ = 0.4941, p=0.6136 by two-way ANOVA, respectively; Figure 4B). Females had higher locomotion than males, with greater average speed and distance traveled during the NOR habituation phase in the open field arena. Male and female SHR treated with eplerenone had decreased speed compared to normotensive controls, and distance traveled was lower in male SHR treated with eplerenone (Figure 4A and 4B; p<0.05 by two-way ANOVA with a post-hoc Tukey test).

**Figure 4.**
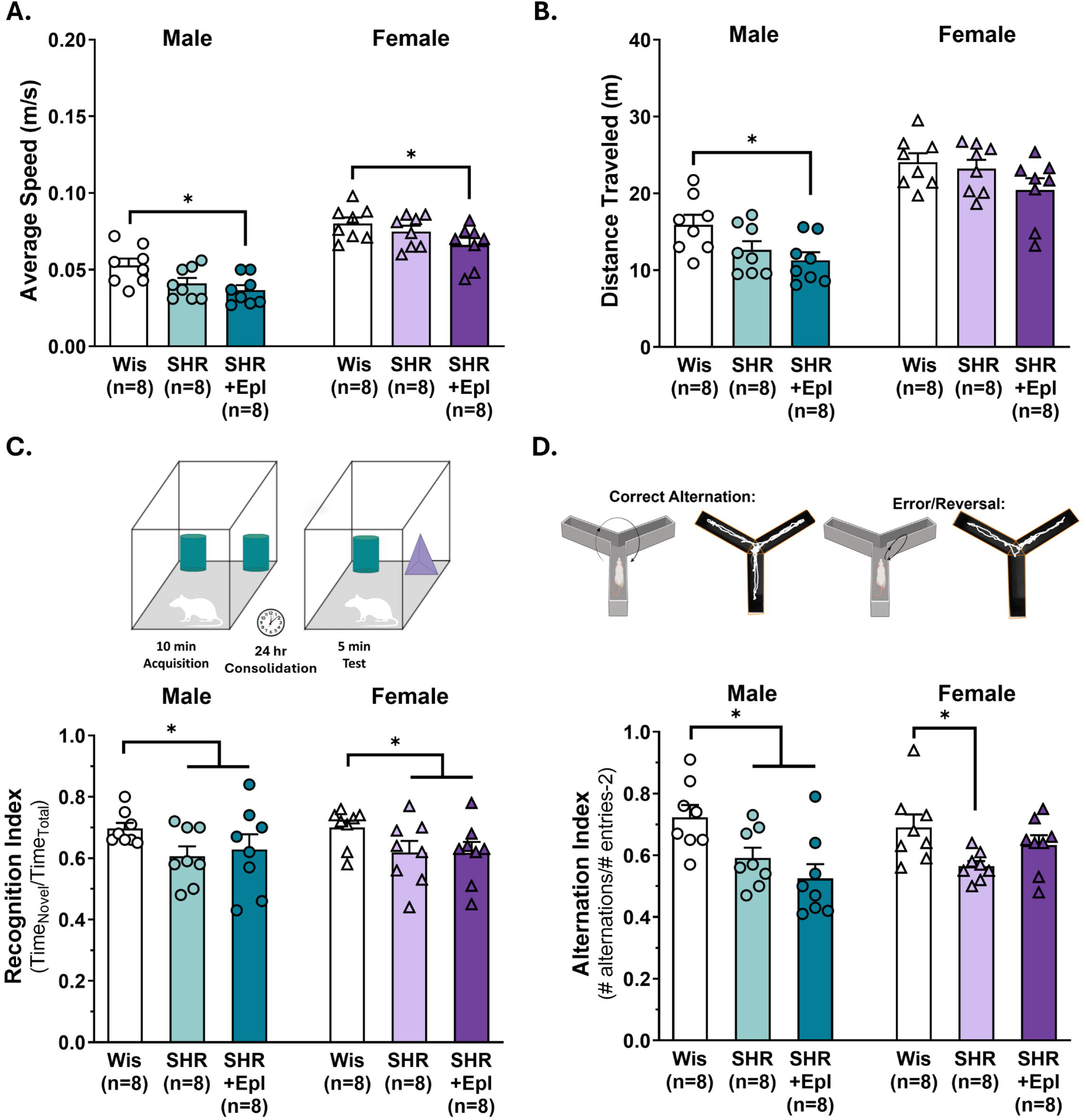
Memory function was impaired during chronic hypertension in a sex- and domain-dependent manner. **A.** Average speed and **B.** distance traveled of male and female normotensive Wistar rats (Wis), spontaneously hypertensive rats (SHR) and SHR treated with mineralocorticoid receptor antagonist eplerenone (SHR+Epl). **C.** Schematic of novel object recognition task used to determine recognition index as a measure of long-term memory function. Recognition index was lower in male and female SHR and SHR+Epl compared to Wis rats. **D.** Schematic of continuous y maze task used to determine alternation index as a measure of spatial working memory. Alternation index was lower during chronic hypertension in males and unaffected by eplerenone treatment. In females, SHR had lower alternation index compared to Wis that was increased by eplerenone treatment. *p<0.05; data are mean ± SEM and analyzed by two-way ANOVA with a post-hoc Tukey test to correct for multiple comparisons.

Figure 4C shows a schematic of the NOR paradigm used to assess long-term memory function, which is largely hippocampal-dependent.^51^ NOR is based upon a rodent’s innate preference for novelty and compares the exploration time of a novel object to that of a familiar object. When given the opportunity, a rat with intact memory function will spend the majority of its time with the novel object (recognition index > 0.50). If memory is impaired and the rat is unable to recognize an object as “familiar”, time is split more evenly between the “novel” and “familiar” objects (recognition index closer to 0.50). Despite differences in locomotion, all rats spent a sufficient amount of time (> 30 seconds) investigating the identical objects during the acquisition phase of NOR for memory encoding. There was a main effect of condition on long-term memory function (F_2, 42_ = 3.726, p=0.0324 by two-way ANOVA; Figure 4C), but there was not an effect of sex and no condition-by-sex interaction (F_1, 42_ = 0.004, p=0.9527; F_2, 42_ = 0.06169, p=0.9403 by two-way ANOVA, respectively). Compared to normotensive rats, object recognition memory was lower in male and female SHR regardless of eplerenone treatment, although there were no statistical differences between groups in the post hoc Tukey test, likely due to within-group variability (Figure 4C).

Spatial working memory largely involves mPFC, and this was tested using a continuous y maze task, as illustrated in Figure 4D.^52, 53^ If working memory is intact, a rat will alternate between maze arms – move from arm A to B to C – without making a reversal entry, which is considered an error. An alternation index was calculated as a measure of spatial working memory, with a higher index indicating better working memory performance. Figure 4D shows a schematic of the task and representative traces of strong memory performance containing arm alternations and poor performance of arm reversals. There was a main effect of condition on spatial working memory (F_2, 42_ = 8.323, p=0.00009 by two-way ANOVA; Figure 4D) but no effect of sex and no interaction between condition and sex (F_1, 42_ = 0.3012, p=0.586; F_2, 42_ = 2.445, p=0.099 by two-way ANOVA, respectively). Alternation index was lower in male and female SHR compared to Wistar rats, which was rescued by eplerenone treatment in females but not males (p<0.05 by two-way ANOVA with post hoc Tukey test). Together, these findings suggest chronic hypertension impaired multiple types of memory function in both sexes, and blockade of MR activity during hypertension selectively improved spatial working memory in females.

Multi-linear regression analyses with Pearson’s correlations were used to generate correlation matrices for blood pressure, hemodynamics in CA1 of hippocampus and object memory (Figure 5A-B) and blood pressure, hemodynamics in mPFC and spatial working memory (Figure 5C-D) in both sexes. In CA1 of males, blood flow was positively correlated with recognition index (p=0.034 by Pearson’s correlation); however, there was no relationship between blood pressure or CA1 PbtO_2_ with long-term memory function (p=0.531, p=0.958, respectively; Figure 5A). CA1 blood flow was negatively correlated with blood pressure (p<0.0001) and positively correlated with PbtO_2_ (p=0.015). There was a negative correlation between blood pressure and CA1 PbtO_2_ (p=0.019 by Pearson’s correlation). In females, CA1 PbtO_2_ positively correlated with recognition index (p=0.022, p=0.305, p=0.121 by Pearson’s correlation, respectively; Figure 5B). There was a negative correlation between blood pressure and blood flow (p<0.0001) and a positive correlation between CA1 blood flow and PbtO_2_ (p=0.002). Blood pressure and CA1 PbtO_2_ were negatively correlated (p<0.0001 by Pearson’s correlation).

**Figure 5.**
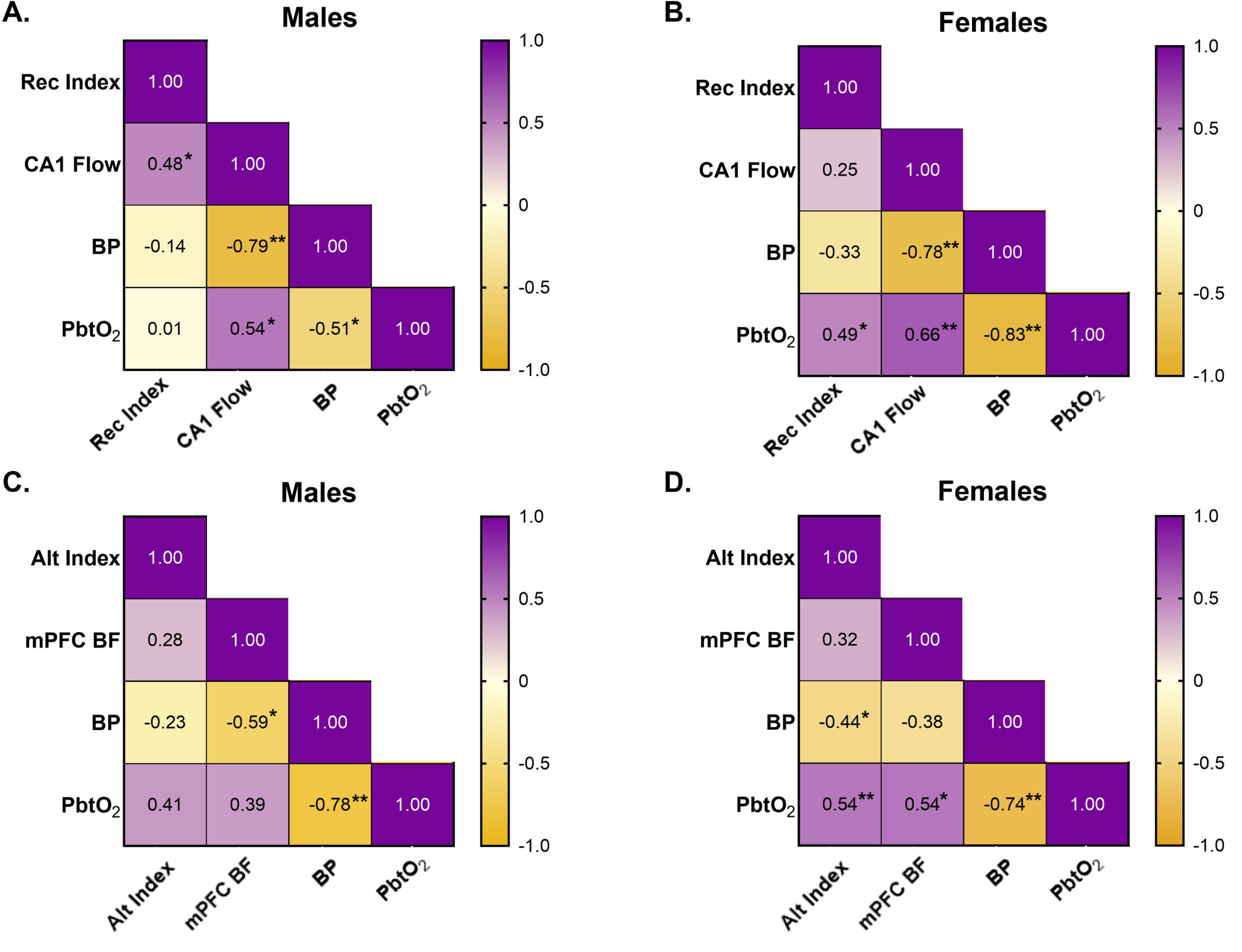
Relationships between cerebral hemodynamics and memory function. A-B. Correlation matrices demonstrating the relationship between long-term memory, as assessed via object recognition index (Rec Index), CA1 blood flow (BF), blood pressure (BP), and partial pressure of brain tissue oxygen (PbtO_2_) in male and female rats. **C-D.** Correlation matrices demonstrating the relationship between spatial working memory, as assessed using an alternation index (Alt Index), medial prefrontal cortex (mPFC) BF, BP, and PbtO_2_ in male and female rats. *p<0.05, **p<0.01; n=19-22/sex; data are analyzed using multi-linear regression analyses with Pearson’s correlations, r^2^ values are reported within the matrix for each comparison.

In regard to spatial working memory, there was no clear relationship between mPFC blood flow or blood pressure with alternation index in males (p=0.239, p=0.283 by Pearson’s correlation, respectively). However, mPFC PbtO_2_ trended towards being positively correlated with spatial working memory, although this did not reach statistical significance (p=0.068; Figure 5C). mPFC blood flow was negatively correlated with blood pressure (p=0.006) and positively correlated with PbtO2, although this did not reach statistical significance (p=0.085). Similarly to CA1, mPFC PbtO2 was negatively correlated with blood pressure (p<0.0001 by Pearson’s correlation; Figure 5C). In females, spatial working memory had no clear relationship with mPFC blood flow (p=0.176), had an inverse relationship with blood pressure (p=0.036), and was positively correlated with mPFC PbtO2 (p=0.008 by Pearson’s correlation; Figure 5D). There was a nonsignificant inverse relationship between mPFC blood flow and blood pressure (p=0.106), and tissue oxygenation was positively correlated with blood flow (p=0.016) and negatively correlated with blood pressure (p<0.0001 by Pearson’s correlation; Figure 5D). Thus, blood pressure was consistently correlated with hypoperfusion in both hippocampus and mPFC in both sexes, and there were stronger relationships between memory, blood pressure, blood flow and tissue oxygenation in females than in males.

## Discussion

There are currently no treatments for VCID. Understanding mechanisms by which modifiable risk factors impact blood flow in cognition-centric brain regions is critical to determine novel therapeutic targets to prevent or delay VCID. Additionally, advancing our understanding of sex-specific mechanisms of hypertension-induced cerebral hypoperfusion and cognitive dysfunction is of utmost importance. Females represent ∼ 50% of the population, develop hypertension in mid-life and have greater longevity than men, yet have largely been excluded from basic cerebrovascular and neuroscience research.^54, 55^ For the first time that we are aware of, we investigated how chronic hypertension disrupted cerebral hemodynamics in both sexes, including the role of aldosterone, and whether this differed by brain region or memory domain to contribute to VCID. We found that chronic hypertension impaired hemodynamics in a sex- and region-dependent manner, with more pervasive perfusion deficits present in females than males. We determined a female-specific role of MR signaling in hypertension-induced cerebral hypoperfusion and memory impairment. This study advances our understanding of mechanisms by which chronic hypertension compromises brain blood flow, highlights the importance of blood pressure control in mid-adulthood, and that efficacy of VCID interventions, specifically targeting aldosterone depend upon sex.

Chronic hypertension has no overt symptoms; it is considered a “silent killer” as it often goes undiagnosed. Hypertension-induced cerebrovascular dysfunction accelerates age-related cerebral hypoperfusion, which is a primary mechanism of neurodegeneration, brain atrophy and VCID.^21, 22^ In the current study, we report that chronic hypertension correlated with lower brain blood flow and impaired long-term and spatial working memory in both sexes; however, there was a differential effect of perfusion based upon brain region and sex. In the dorsal hippocampus, chronic hypertension decreased CA1 perfusion in both sexes, but to a greater extent in males than females. In the mPFC, there were perfusion deficits only in hypertensive females, with little impact in males, although the data were more variable. Importantly, PbtO_2_ was substantially reduced in both brain regions in SHR of both sexes, including mPFC in males even in the absence of perfusion deficits. This demonstrates that chronic hypertension caused wide-spread hemodynamic disruption impacting both anterior (via carotid system) and posterior (via vertebrobasilar system) cerebral circulation territories. However, blood flow and metabolism remained tightly coupled under chloral hydrate anesthesia. While perfusion and PbtO_2_ were decreased in CA1 and mPFC during hypertension, tissue oxygenation did not drop to levels considered to be detrimentally hypoxic (e.g., less than ∼ 20 mmHg), assuming neuronal hypoxic/ischemic tolerance is similar between normotensive and hypertensive states.^56–58^ However, anesthesia reduces neuronal activity and CMRO_2_ and, based upon the Fick Principle, would be reflected as higher PbtO_2_, suggesting the values in the current study are an overestimation compared to in the unanesthetized state. As PbtO_2_ levels were determined to be between ∼ 20-22 mmHg in SHR, tissue hypoxia may be present in both brain regions in mid-adulthood in awake rats and contribute to impaired long-term and spatial working memory in SHR. We have previously shown that chronic hypertension in males impaired hippocampal vascular function that reduced hippocampal blood flow and impaired memory as a function of age.^16^ Since the current study investigated cerebral hemodynamics during in mid-adulthood, it was not determined whether progressive perfusion deficits result in hypoxia/ischemia and VCID at older ages.

Primary aldosteronism is present in up to 20% of people with uncontrolled, drug-resistant hypertension.^25^ MRs are expressed in cerebrovascular endothelium and vascular smooth muscle, and activation causes oxidative stress, impaired endothelial-dependent vasodilation and vasoconstriction as well as hypertension-induced structural remodeling.^59–62^ As the most powerful determinant of brain blood flow is lumen diameter of arteries and arterioles, even relatively small decreases in lumen diameter can drastically increase vascular resistance and reduce perfusion.^63^ Here, we show that MR antagonism and had a moderate anti-hypertensive effect and reversed hypertension-induced hypoperfusion in dorsal hippocampus and mPFC exclusively in the female sex. Interestingly, MR antagonism improved spatial working memory in females that is largely mPFC-dependent but did not rescue long-term memory function, despite restoring hemodynamics in dorsal CA1. In contrast to previous studies, we report no role for MRs in hypertension-induced hypoperfusion or memory dysfunction in males, potentially due to differences in hypertensive models (i.e., genetic, stroke-prone, angiotensin II-infusion-mediated) or specific behavioral paradigms.^26, 39^ Regardless, these findings suggest an MR-dependent mechanism of VCID in females, that resilience of memory function differs by domain during chronic hypertension, and that early intervention may be necessary to prevent any degree of hypoperfusion in CA1 to protect hippocampal function and VCID.

Previous studies have reported that MR antagonism in multiple models of hypertension, including stroke-prone SHR, promoted increased perfusion in males, but not females, by improving endothelial function and preventing inward structural remodeling of pial arteries and penetrating arterioles independently of decreasing blood pressure.^26–28, 37, 38^ These findings are in direct contrast to those of the current study that demonstrate a female-specific role of MR signaling in cerebral hypoperfusion during hypertension, despite dose and duration of eplerenone treatments being similar. There are a few potential reasons for these conflicting findings: 1. Previous studies used *ex vivo* pressure myography to assess structural and functional properties of cerebral arteries and arterioles that may not reflect structure/function changes to the vascular network *in vivo* to cause changes in blood flow. 2. Differences may be due to MR expression differing by vascular territory/brain region. Previous studies focused predominantly on the middle cerebral artery territory while the current study investigated hippocampus and mPFC which are in posterior cerebral artery and anterior cerebral artery territories, respectively. Interestingly, despite MR antagonism having no effect in hypertensive females, it was determined that MR expression was higher in pial arteries from female compared to male stroke-prone SHR.^37^ If female SHR have increased MR expression in arterioles perfusing hippocampus and mPFC, that could contribute to the sex-specific role of MRs in the current study. Reliable aldosterone values in plasma were unable to be determined in the current study due to anesthesia- and surgical stress-induced changes in circulating levels.^64^ Alternatively, vascular MR activation causes vasoconstriction via involvement of L-type voltage-gated calcium channels (L-VGCC).^38^ Whether there are sex-specific increases in MR or L-VCGG expression or calcium sensitivity of vascular smooth muscle in CA1 and mPFC remain unclear, but could result in more robust vasoconstriction upon MR activation and contribute to hypoperfusion in females. Regardless, our findings demonstrate that the role for MRs in reductions in brain blood flow during hypertension depends upon sex, and that treatments targeting aldosterone may be more effective at protecting against VCID in females.

During healthy aging, cerebral perfusion decreases and contributes to age-related cognitive decline.^18, 19^ In the current study, we found absolute basal blood flow was similar in dorsal CA1 and mPFC in mid-adulthood of healthy females and males. Despite these similarities in perfusion, PbtO_2_ was consistently higher in both brain regions in females, and there was a significant positive relationship between PbtO_2_ and long-term and spatial working memory in females but not males. CMRO_2_ is the critical link between blood flow and PbtO_2_. As calculated in the current study, increased PbtO_2_ in CA1 and mPFC in females was associated with lower CMRO_2_. It remains unclear whether differences in neuronal metabolism exist in the unanesthetized state or if there was greater oxygen diffusion into these regions in the female sex due to vascular network differences (e.g., capillary density). The findings from the current study suggest that sex differences in PbtO2 may be protective of brain health and delay age-related cognitive decline in healthy females.

Primary aldosteronism in women often correlates with neuropsychiatric symptoms, such as anxiety and depression that involve the ventral hippocampus (anterior in humans).^65^ The current study focused on hemodynamics of dorsal hippocampus due to its primary role in memory, and anxiety-like behaviors were not investigated. However, we speculate that if perfusion deficits extend to ventral hippocampus, it could contribute to the propensity of neuropsychiatric symptoms in women with high aldosterone and increased MR activity. Additionally, stress hormones (e.g., corticosterone/cortisol) are also ligands of MRs and MR antagonism blunts the stress response, reduces neuroinflammation and protects hippocampal neurons.^65^ Thus, it is possible that MR antagonism in the current study impacted multiple cell types. Future studies are needed to distinguish between protective actions of eplerenone on neuronal and arteriolar function, and within the neurovascular unit (e.g., pericytes, astrocytes).

Overall, the findings in the current study support a protective role of MR antagonism on cerebral hemodynamics and memory function in females but not males. Longer treatment duration or combination treatments with traditional anti-hypertensive medications (e.g., captopril) may be necessary to provide a similar level of neurovascular protection in the male sex. This study demonstrated that, in females, hippocampal-dependent memory function was more vulnerable to hypertension-induced perfusion deficits. In contrast to improvement of spatial working memory, long-term memory function was unable to be rescued despite restoration of hemodynamics in CA1. Such domain-specific resilience highlights the critical need for blood pressure control in mid-adulthood in both sexes, and that MR antagonism may be more effective at mitigating VCID in the female sex.

## Acknowledgements

None.

## Data Availability

All datasets generated and/or analyzed during the current study are publicly available within this manuscript and deposited in the Harvard Dataverse repository (https://doi.org/10.7910/DVN/LCZRNS)

## Authorship Statement

GDM contributed via investigation, formal analysis, visualization, writing-original draft, writing-review & editing. GAB, DC, JB, EDS, MEW, EJB and JL contributed via investigation, formal analysis, writing-review & editing. ACC contributed conceptualization, methodology, validation, formal analysis, investigation, resources, data curation, visualization, writing – original draft and writing – review & editing.

## Consent Statement

Consent was not necessary.

